# Intestinal epithelium-derived high density lipoprotein restrains liver injury via the portal vein

**DOI:** 10.1101/2020.10.01.322925

**Authors:** Yong-Hyun Han, Emily J. Onufer, Li-Hao Huang, Rafael S. Czepielewski, Brad W. Warner, Gwendalyn J. Randolph

## Abstract

Assembly of high density lipoprotein (HDL) requires apoA1 and the cholesterol transporter ABCA1. Although the liver generates most HDL in blood, HDL synthesis also occurs in the small intestine. However, distinct functions for intestinal HDL are unknown. Here we show that HDL in the portal vein, which connects intestine to liver, derivesd mainly from intestine and potently neutralizes lipopolysaccharide (LPS). In a mouse model of short bowel syndrome where liver inflammation and fibrosis was driven by the LPS receptor TLR4, loss of intestine-derived HDL worsened liver injury, whereas liver status was improved by therapeutics that elevated and depended upon intestinal HDL production. Thus, protection of the liver from injury in response to gut-derived signals like LPS is a major function of intestinally synthesized HDL.

---

The portal vein collects venous drainage from the intestine, carrying nutrients and metabolites of host and microbiome origin directly to the liver as its major blood supply (*1*). Through this route, components of the microbiome may drive liver injury, characterized by steatohepatitis, cholestasis, and fibrosis (*2, 3*). Indeed, intestine-derived lipopolysaccharide (LPS) from Gram-negative bacteria triggers inflammatory and fibrotic pathways in the liver after binding TLR4 (*3, 4*). TLR4 engagement may also underlie intestinal failure-associated liver disease that results from surgical resection of large portions of the small intestine as a complication of necrotizing enterocolitis, Crohn’s disease, cancer, or trauma (*5*–*7*).

Given the links between TLR4 driven liver injury and the general abundance of LPS in the intestine, we wondered whether the host may have evolved an unidentified mechanism to limit LPS-mediated liver injury via the gut-portal axis. In this regard, we were led to consider that high density lipoprotein (HDL) might have an overlooked role in host protection of the liver via its long acknowledged potential to neutralize LPS (*8*–*10*). Indeed, it is unclear why HDL is additionally synthesized by the intestine rather than solely by the liver. HDL-cholesterol (HDL-C) is the smallest lipoprotein particle in the blood and is best known for its ability to collect cholesterol peripherally and deliver it to the liver for ultimate disposal in the bile, a process known as reverse cholesterol transport. Only two tissues produce apoA1, the core protein component of HDL-C: the liver and the small intestine (*11*). A distinct function for intestinal HDL had seemed unlikely as mice whose intestinal epithelial cells selectively delete the gene encoding the cholesterol transporter ABCA1, which facilitates the formation of nascent HDL, manifest only a modest (25-30%) reduction in plasma HDL-C; in constrast, there is a great magnitude of reduction observed after liver-specific loss of ABCA1 (*12*), leaving investigators to consider the intestine as simply a second source of HDL-C.

An obstacle to considering a possible link between intestinal HDL and LPS actions on the liver is the paucity of knowledge concerning a pathway for intestine-derived HDL delivery to the liver. HDL typically transports from tissues through lymphatic vessels (*13*–*15*), which do not converge directly to the liver (*16*). On the other hand, an earlier study was unable to demonstrate that intestinally produced HDL-C entered lymphatics, however, an alternative entry, via the portal vein, was not examined (*17*). Here, we show that, remarkably, not only is intestinally generated HDL-C transported primarily into the portal circulation, but nearly all HDL-C found in the portal blood is derived from the intestine. Intestinal epithelial cells produced small HDL particles (HDL_3_) (*18*) with potent anti-inflammatory and LPS-neutralizing properties that, rather than entering lymphatics, directly entered portal blood. After small bowel resection in mice, intestinally derived HDL indeed had a key role in protecting the liver from the onset of inflammatory and fibrotic damage. By contrast, elevation of intestinal HDL using a therapeutic approach protected against liver failure after small bowel resection. Thus, targeting HDL in the intestine may have novel applications to promote liver health.

## Results

### Intestine-derived HDL is the main source of HDL in portal blood

We first evaluated apoA1 levels in portal blood compared with systemic blood (from the inferior vena cava) and found that, relative to albumin, apoA1 was ∼40% lower in portal plasma than systemic plasma in both humans and mice (Fig. 1A). This reduced concentration was accompanied by an unexpectedly diminished recirculation of systemic HDL into portal blood. That is, when we used photoactivatable apoA1 knockin mice (PGA1^KI/+^), that express from the mouse apoA1 gene locus a version of human apoA1 with a photoactivable GFP tag at the N-terminus (*15*), and photoactivated apoA1 in the skin of this knock-in strain, we later observed skin-tagged HDL in the systemic blood and mesenteric lymph, but it was very low in the portal blood (Fig. 1B). Its presence in mesenteric lymph to a concentration that approached that in the systemic circulation (Fig. 1B) suggested that systemic HDL escaped the blood circulation prior to entering the portal venous blood to enter the gut interstitium and was reabsorbed by the lymphatics, reminiscent of some i.v. injected protein antigens (*19*) (Fig. 1B).

**Figure 1.**
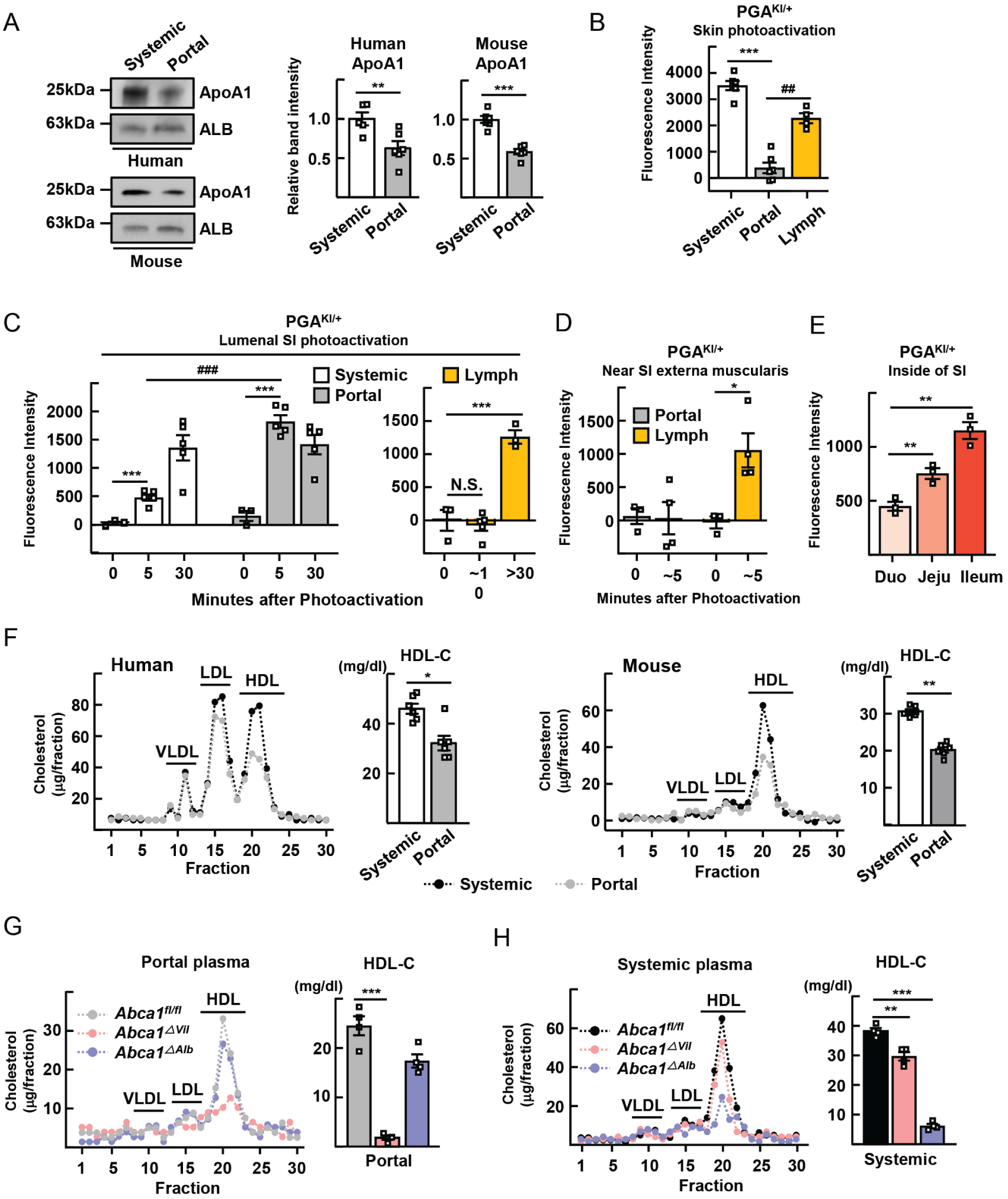
Intestinal HDL transits through portal vein and accounts for most HDL in the portal blood. (**A**) Immunoblot for apoA1 in paired systemic and portal from humans or mice. (**B**) Human HDL was IV injected into WT mice and assayed 2 h later (left). Skin of PGA1^KI/+^ mice was photoactivated; plasma and lymph fluorescence was measured 2 h later (right). (**C and D**) The lumen (**C**) or exterior (**D**) of the small intestinal ileum (SI) of PGA1^KI/+^ mice was photoactivated; plasma and mesenteric lymph fluorescence was assessed 5-10 or 30 min later. (**E**) The lumen of duodenum, jejunum, or ileum of PGA1^KI/+^ mice was photoactivated; portal plasma fluorescence assessed 5 min. later. (**F**) Lipoprotein profile of systemic and portal plasma from human (n=6) or mouse (n=6). (**G and H**) Lipoprotein profiles and HDL-C quantification of portal (**G**) or systemic (**H**) plasma from *Abca1*^*fl/fl*^, *Abca1*^Δ *Vil*^, and *Abca1*^Δ *Alb*^ mice. Data are plotted as mean ± SEM, male and female mice combined, with each symbol on the bar graphs representing a single mouse. *P < 0.05; **P < 0.01; ***P < 0.001; ^##^P < 0.01; ^###^P < 0.001.

Given the strikingly diminished presence of systemic HDL in the portal blood in PGA1^KI/+^ mice, we wondered if the intestine might be a key source of HDL in the portal vein. To examine this, we again utilized the PGA1^KI/+^ mouse by slipping a small 405-nm laser into the lumen of the small intestine to photoactivate the intestinal epithelium directly. Within 5 minutes, strong fluorescence was detected in the portal blood that at this early time point was more than 3-fold higher than the systemic circulation (Fig. 1C), with equilibration between the portal and systemic compartments after 30 minutes (Fig. 1C). At 30 minutes but not 5 minutes after epithelial photoactivation, the mesenteric lymph draining from the ileum was fluorescent when activation occurred at the intestinal lumen. If, on the other hand, the activation was carried out by photoactivation of the exterior of the intestine, outside the muscularis, the lymph but not the portal vein rapidly acquired strong fluorescence (Fig. 1D). These data indicate that HDL tagged at the intestinal epithelial interface first enters the portal vein and is not observed in lymph until the portal vein has brought blood to the liver and it mixes with systemic blood, such that the passage of systemic blood in intestinal layers like the muscularis allows for HDL recirculation into lymph.

The ileum was the major site of HDL formation in the intestine based on studies wherein we photoactivated duodenum, jejunum, or ileum (Fig. 1E). Just as apoA1 was overall lower in the portal blood, HDL-C was also quantitatively reduced in portal plasma compared with systemic plasma in both humans and mice (Fig. 1F). We separated the different classes of lipoproteins via fast protein liquid chromatography (FPLC) and found that HDL-C in particular was decreased by >90% in portal blood of intestine-specific ABCA1 knockout mice (*Villin*^Cre^-*Abca1*^fl/fl^; *Abca1*^Δ *Vil*^) (Fig. 1G). This dramatic decrease was restricted to the portal blood, as we observed only ∼25% decrease in HDL-C from systemic blood of the same *Abca1*^Δ *Vil*^ mice (Fig. 1H), as previously reported (*12*). In contrast, liver-specific ABCA1 knockout mice (*Abca1*^Δ *Alb*^) had a marked reduction of HDL-C in systemic blood, but not portal blood (Fig. 1G, H). Collectively, these data establish two distinct, differentially controlled blood compartments for HDL, one in the portal venous blood governed by intestinal production of HDL, and the other in systemic vessels outside of the portal venous system, governed by liver production of HDL.

### Portal blood HDL is small-sized and strongly neutralizes LPS-mediated inflammatory responses

Portal venous HDL was relatively small in size (∼8 nm) in both human (Fig. 2A) and mouse (Fig. 2B) as assessed by apoA1 immunoblotting under nondenaturing conditions (Fig. 2A, B, left) and by transmission electron microscopy (Fig. 2A, B, right). HDL particles are classified based on physicochemical properties, including size. Specifically, large-sized HDL_2_ and small-sized HDL_3_ particles are composed respectively of distinct accessory proteins, with paraoxonase 1 (PON1) enriched in HDL_3_ particles, and apoB in HDL_2_ particles (*18*). Besides their small size, portal venous HDL was enriched in PON1 and low in ApoB and thus best classified as HDL_3_ (Fig. 2C). Altogether, these data indicate that portal venous HDL is principally HDL_3_.

**Figure 2.**
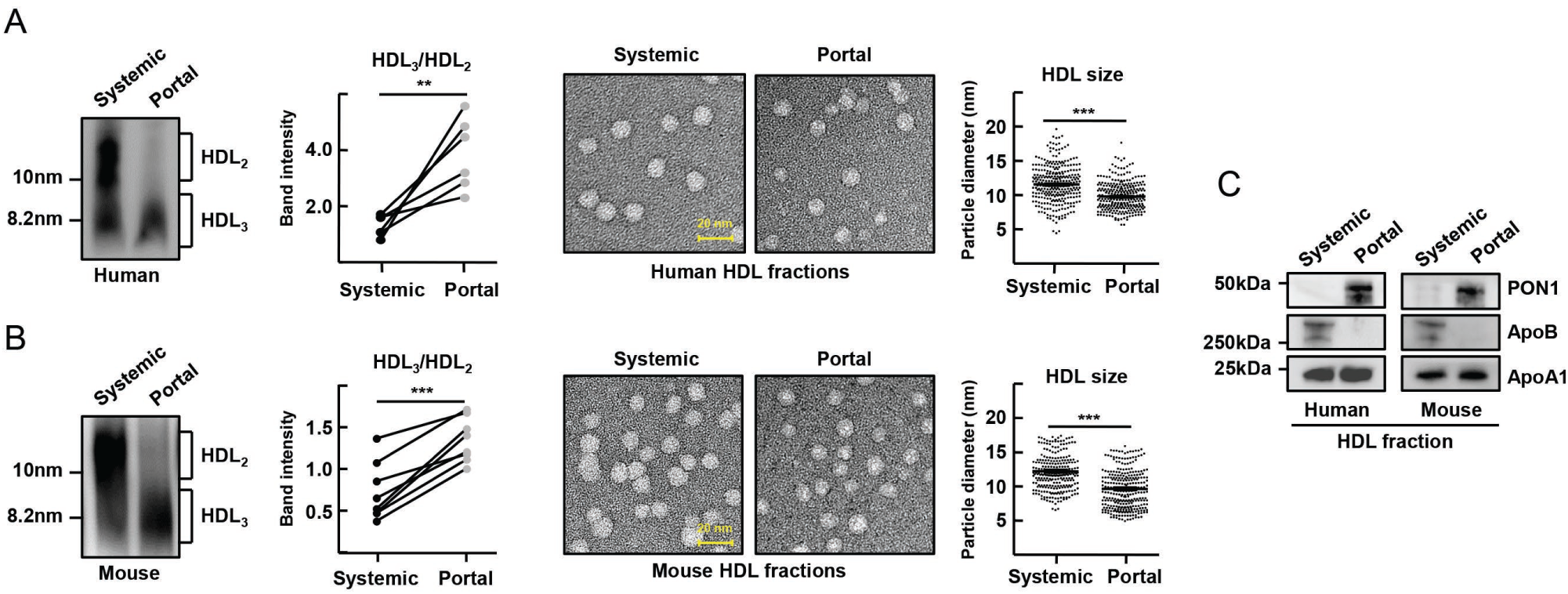
Portal vein HDL is enriched in small HDL_3_ particles. (**A and B**) Immunoblots for apoA1 from nondenaturing gels using human (**A**) and mouse (**B**) systemic versus portal plasma HDL. Representative gels and plots show HDL_3_/HDL_2_ in paired systemic and portal plasma samples (left). Representative electron microscopy images of negative-stained HDL fractions from systemic and portal plasma. HDL particle diameter; each dot represents a single measured particle (right). (**C**) PON1, apoB, and apoA1 in HDL fractions analyzed by immunoblot. Error bars in plots show mean and SEM, combining data from male and female mice. **P < 0.01; ***P < 0.001.

Considering the known capacity of HDL to neutralize LPS (*8, 9*), we wondered if a relevant function for HDL transiting the gut-liver axis through the portal vein might be the neutralization of portal blood LPS, likely arising from the microbiome-rich ileum, as such neutralization might dampen LPS signaling and inflammatory signals in the liver. To test how HDL affected liver macrophage responses to LPS, we isolated primary liver Kupffer cells expressing high Clec4F, TIMD4, and F4/80 (*20, 21*) (Fig. S1) from WT or TLR4^−/−^ mice. Then we stimulated the macrophages with LPS mixed with, or not, HDL from the portal vein versus peripheral venous blood of humans (Fig. 3A) or mice (Fig. 3B). Portal venous-derived HDL fractions obtained from either species more effectively neutralized LPS-induced pro-inflammatory responses in Kupffer cells than HDL from systemic blood or no HDL at all, as assessed by analysis of mRNAs for inflammatory genes, production of CCL2, or iNOS expression (Fig. 3A, B). Whole portal plasma obtained from *Abca1*^Δ *Vil*^ mice less effectively protected against induction of inflammation as compared with portal plasma from WT or *Abca1*^Δ *Alb*^ mice (Fig. 3C), illustrating that intestinal HDL was a prominent overall anti-inflammatory mediator in the portal vein.

**Figure 3.**
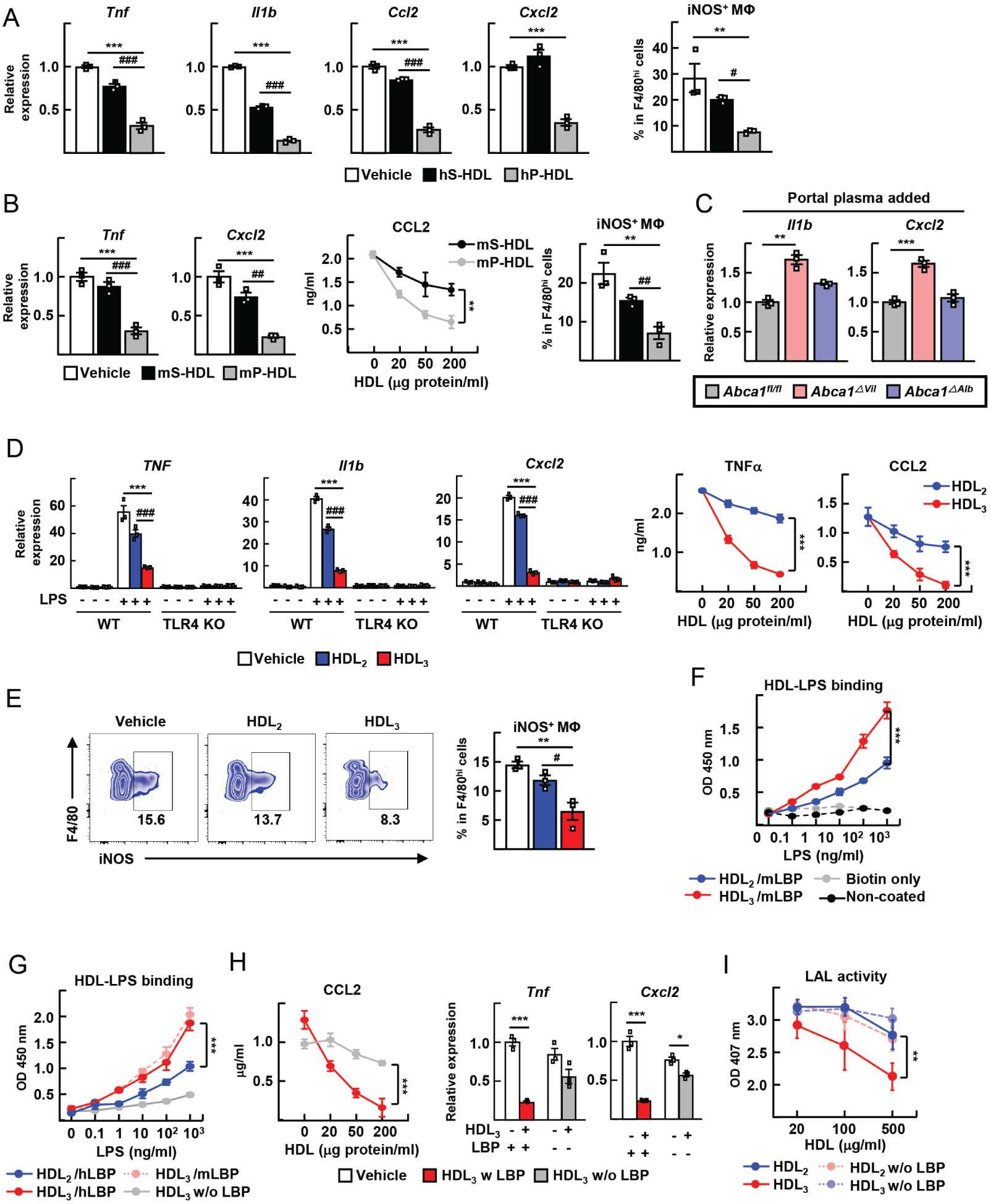
Portal blood HDL_3_ has stronger LBP-mediated anti-inflamatory action on liver macrophages. (**A, B**) LPS-treated liver macrophages were incubated with 100 μg/ml HDL obtained from human systemic plasma (hS-HDL) or human portal plasma (hP-HDL) prior to preparation of mRNA, FACS phenotyping (iNOS^+^ F4/80^hi^ macrophages), or CCL2 ELISA. (**C**) LPS-treated liver macrophages incubated with 5% portal vein-derived plasma obtained from *Abca1*^*fl/fl*^, *Abca1*^Δ *Vil*^, and *Abca1*^Δ *Alb*^ mice. (**D and E**) WT and TLR4^−/−^ liver macrophages were incubated in serum-free LBP-containing medium with or without 20 ng/ml LPS and 100 μg/ml HDL_2_, HDL_3_ before analysis of mRNA, FACS phenotyping (iNOS^+^ F4/80^hi^ cells), or TNFα and CCL2 ELISA. (**F and G**) ELISA to assess binding of HDL and biotin-LPS with or without LBP added. (**H**) LPS-treated liver macrophages were incubated with 100 μg/ml HDL_3_ in the presence or absence of 1 μg/ml LBP. Production of CCL2 measured by ELISA (left); mRNA transcripts of inflammatory genes measured by qRT-PCR (right). (**I**) Endotoxin LAL activity assessed after 0.5 EU/ml *E. coli* LPS was pre-incubated with different concentrations of HDL with or without 1 μg/ml LBP. All data are plotted as mean ± SEM using a combination of male and female mice, with n=3-5 per time point. Each symbol represents data obtained using a different mouse. *P < 0.05; **P < 0.01; ***P < 0.001; ^#^P < 0.05; ^##^P < 0.01; ^###^P < 0.001.

To consider whether the stronger LPS neutralizing activity of portal venous HDL could be attributed to HDL_3_ or was related to some as yet undefined property of portal venous HDL, we repeated the experiments using human HDL_2_ or HDL_3_ from peripheral blood. Indeed, HDL_3_ strongly suppressed LPS-induced transcriptional activation of inflammatory genes compared with HDL_2_ (Fig. 3D) and suppressed iNOS induction (Fig. 3E), with all inflammatory activity being dependent upon Kupffer cell expression of TLR4 (Fig. 3D). HDL_3_ also decreased TNFα and CCL2 secretion in LPS-stimulated Kupffer cells in a dose-dependent manner (Fig. 3D). These data indicate that HDL_3_, the primary HDL subclass found in human or mouse portal blood, potently suppresses LPS-induced proinflammatory activation of liver macrophages.

Next, we quantified the efficiency of binding between LPS and HDL using a sandwich ELISA, modifying a previous approach (*22*). HDL was immobilized on plates and subsequent binding of biotinylated-LPS was tracked with a streptavidin detection reagent. Human HDL_3_ more robustly associated with LPS compared with HDL_2_ in the presence of murine LPS-binding protein (LBP) (Fig. 3F), a plasma protein that, like PLTP and CETP, is in the TULIP family of lipid binding and transport proteins and mediates delivery of LPS to CD14 that then facilitates signaling through TLR4 (*23*). LBP associates with HDL in the presence of LPS (*22, 24*). Human LBP incubated with HDL_2_ or HDL_3_ still showed preferential binding of LPS for HDL_3_, which bound LPS equally well regardless of the presence of human or mouse LBP, though LBP from at least one of these sources was required for binding (Fig. 3G). LBP also strongly promoted the anti-inflammatory effects of HDL_3_ on Kupffer cells (Fig. 3H). When we employed the classical Limulus assay to assess LPS activity, the presence of LBP + HDL_3_ clearly neutralized LPS activity in the assay, whereas LBP + HDL_2_ did not lead to neutralized activity (Fig. 3I), despite some binding (Fig. 3F). Thus, HDL_3_ particles in particular potently bind and neutralize LPS, leading to reduced inflammatory activation of liver macrophages.

### The LPS receptor TLR4 on bone marrow cells drives liver injury in response to small bowel resection

Given our findings that intestinally produced HDL entered the portal vein in a form that potently neutralized LPS, we next aimed to determine whether there was a role in vivo for HDL neutralization of LPS in modulating liver inflammation in response to conditions in the gut that might allow more LPS to enter the portal vein. Surgical resection of significant portions of the small intestine of mice has been reported to promote hepatic steoatosis in a TLR4-dependent manner (*25*). Furthermore, this surgical procedure in mice, as in humans, promotes liver fibrosis (*26*). The role of TLR4 in the small bowel resection-induced fibrosis remains unestablished, but might be anticipated based on the role of TLR4 in liver fibrosis in other models (*27*).

First, we established a variety of inflammatory changes in mice that received small bowel resections, wherein we removed 50% or 75% of the small intestine (Fig. 4A), sparing the head of the small intestine and the terminal portion of the ileum, except where indicated. Within 3 months, these resections induced morphological changes in the liver (Fig. 4B), elevated plasma aspartate aminotransferase (AST) in plasma (Fig. 4C), and enhanced infiltration of myeloid cells (F4/80^+^ and S100A9^+^) in the liver (Fig. 4D, Fig. S2A) that was associated with transcriptional induction of inflammatory mediators and upregulation of fibrosis indciators (Fig. S2B and S2C). Resection of 75% of the intestine led to a more severe outcome than 50% resection, as previously observed (*26*). LPS activity in the portal vein was elevated by small bowel resection (Fig. 4E), and we wondered if this was linked to higher intestinal and portal vein permeability that could lead to more LPS escaping the intestine and entering the portal vein (*1*). Indeed, small bowel resection led to increased intestinal permeability, assessed by passage of fluorescein isothiocyanate (FITC)-dextran from the intestine to the systemic circulation (Fig. 4F). Furthermore, mRNA levels of epithelial junction proteins ZO-1 and occludin (Ocln) were reduced (Fig. 4G), while the venous capillaries in intestinal villi expressed elevated plasmalemma vesicle-associated protein 1 (PV1) (Fig. 4H), indicative of more porous fenestrations of the vessels feeding into the portal vein (*28*).

**Figure 4.**
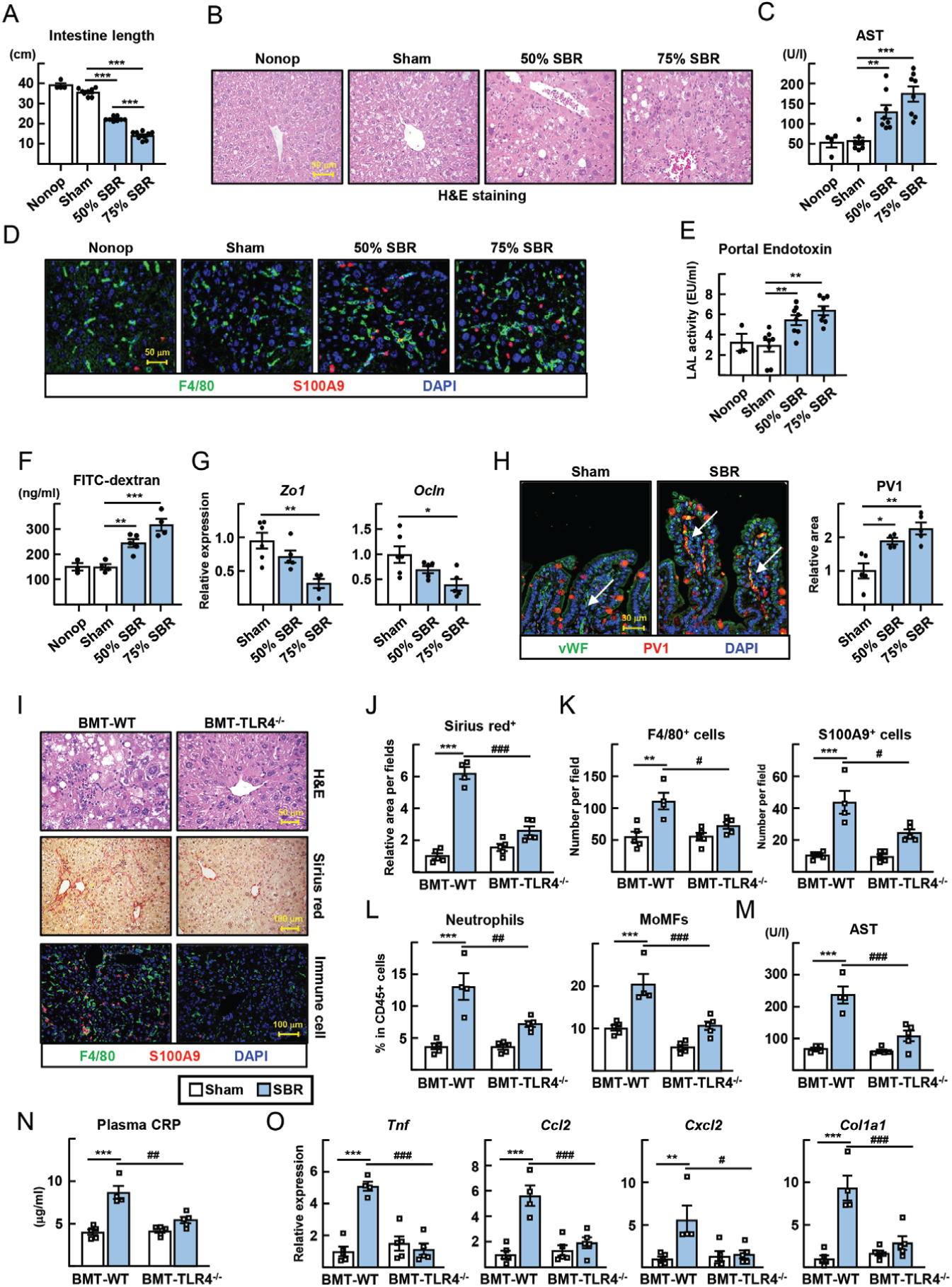
Small bowel resection triggers TLR4-mediated liver inflammation. (**A-H**) Small bowel resection (SBR) operations were conducted on WT mice. Non-operated (Nonop) (n=4), sham (n=8), 50% SBR (n=8), and 75% SBR (n=8) mice were euthanized 12 weeks later. (**A**) The total length of small intestines. (**B**) Representative H&E images of liver sections. (**C**) Plasma AST levels. (**D**) The F4/80^+^ macrophages and S100A9^+^ neutrophils were stained in liver sections. (**E**) LAL endotoxin activity of portal plasma was measured. (**F**) Quantitative analysis of FITC-dextran translocation from intestine to peripheral blood. (**G**) Intestinal mRNA transcripts encoding tight junction proteins analyzed by qRT-PCR. (**H**) Expression of von Willebrand Factor (vWF) (blood vessel) and PV1 was visualized in intestinal sections (left). Note that goblet cell mucin stains with the PV1 antibody, possibly nonspecifically. The relative increase in PV1 staining of vWF^+^ vessels (vessels highlighted by white arrows) after short bowel resection was quantified (right). (**I-O**) Mice receiving bone marrow transplants with WT or TLR4^−/−^ hematopoietic cells received SBR and analyzed 10 weeks later. (**I**) H&E, sirius red, and immunostaining of liver sections. (**J**) The relative sirius red positive area per field (**K**) Numbers of F4/80^+^ macrophages and S100A9^+^ neutrophils per field. (**L**) The proportions of hepatic neutrophils and monocyte-derived macrophages (MoMFs) among CD45^+^ leukocytes. (**M and N**) Plasma AST and CRP levels. (**O**) Hepatic mRNA transcripts of inflammatory genes analyzed by qRT-PCR. All data are plotted as mean ± SEM in male mice, with each symbol on the bar graphs representing a single mouse. *P < 0.05; **P < 0.01; ***P < 0.001; ^#^P < 0.05; ^##^P < 0.01; ^###^P < 0.001.

To determine if TLR4 was required for liver inflammation and collagen deposition after small bowel resection and to explore a possible role of TLR4 expressed by radiosensitive cells like liver macrophages, we transplanted TLR4-deficient bone marrow cells into irradiated wild type mice (BMT-TLR4^−/−^) and compared them after small bowel resection to mice receiving WT bone marrow. Among hematopoietic cells, TLR4 is selectively expressed by macrophages, so any effects of the transplant on liver inflammation after bowel resection would suggest a role for TLR4 on macrophages. Sirius red staining of liver sections indeed revealed reduced fibrosis in the BMT-TLR4^−/−^ group (Fig. 4I-J). Infiltrates of myeloid cells in the liver (Fig. 4K, L) and induction of AST and C-reactive protein (CRP) were also greatly reduced in BMT-TLR4^−/−^ mice receiving small bowel resection (Fig. 4M, N), as were inflammatory mediators (Fig. 4O). Thus, extensive intestinal resection, mimicking a clinical relevant condition, leads to alterations in the gut that heighten permeability in general and LPS translocation to the liver via the portal vein, with TLR4, the receptor for LPS, driving consequent liver injury.

### Disruption of intestine-derived HDL exacerbates liver injury after small bowel resection

These findings set the stage to next address whether HDL modulates TLR4-driven liver injury that follows intestinal resection. Portal venous HDL-C was lowered after small bowel resection (Fig. 5A), possibly a consequence of the loss of bowel mass that might normally contribute to HDL biogenesis. Expression of ABCA1 sharply elevated from proximal to distal small bowel, whereas apoA1 was more uniformly expressed with gradual enhancement in distal regions of the small intestine, overall suggesting that the ileum is the most critical region for HDL production in the bowel (Fig.5B). When we modified the region of the bowel resected to remove the proximal 50% (P-SBR) or distal 50% (D-SBR) of the small intestine, HDL-C in portal blood fell most drastically after the distal resection (Fig. 5C). Fitting with our hypothesis that reduced HDL-C levels in portal blood would lead to less protection of the liver against injury, liver injury and inflammatory markers were greater in the D-SBR group (Fig. 5D-F).

**Figure 5.**
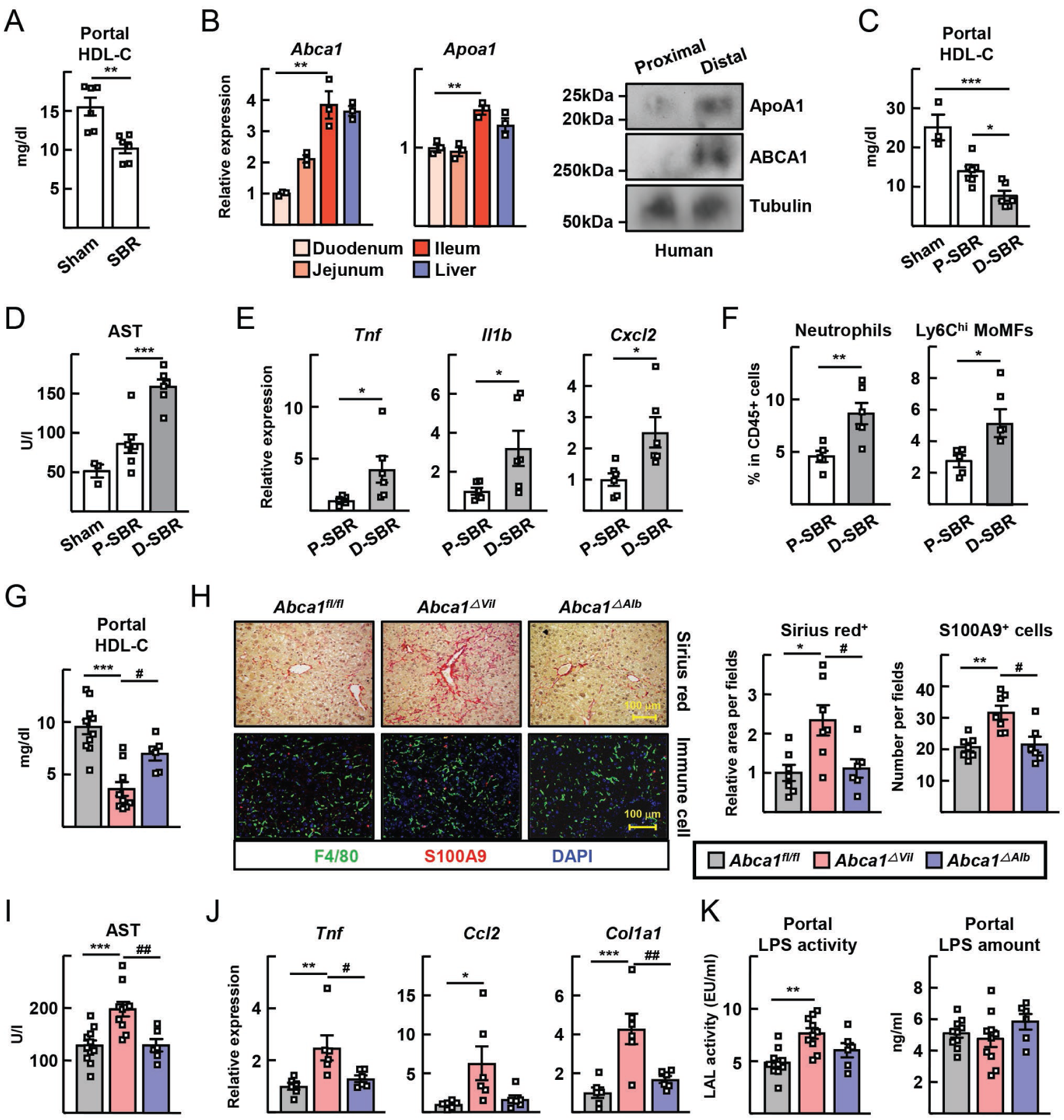
Intestinal disruption of HDL production worsens short bowel resection-induced liver injuries. (**A**) Portal HDL cholesterol levels measured in sham and 75% short bowel resection (SBR) mice. (**B**) mRNA transcripts of *Abca1* and *Apoa1* in mice duodenum, jejunum, ileum and liver were measured by qRT-PCR (left). The protein expression of apoA1 and ABCA1 in human proximal and distal gut were analyzed by immunoblotting (right). (**C-F**) SBR operations were conducted for WT mice. Mice receiving sham (n=3), proximal 50% SBR (P-SBR) (n=7), and distal 50% SBR (D-SBR) (n=6) operations were euthanized 10 weeks later. Portal HDL cholesterol levels (**C**). Plasma AST levels (**D**). Hepatic mRNA transcripts of inflammatory genes (**E**). The proportions of hepatic neutrophils and Ly6C^+^ monocyte-derived macrophages (MoMFs) among CD45^+^ leukocytes (**F**). (**G-K**) 75% SBR operations were performed for *Abca1*^*fl/fl*^ (n=10), *Abca1*^Δ *Vil*^ (n=10), and *Abca1*^Δ *Alb*^ (n=6) mice and euthanized 8 weeks later. Portal HDL cholesterol levels (**G**). Representative images of sirius red staining of liver sections (left) and relative area per field (right). The F4/80^+^ macrophages and S100A9^+^ neutrophils were visualized (left); cell numbers per field (right) (**H**). Plasma AST (**I**). Hepatic mRNA transcripts of inflammatory genes (**J**). LPS activity by LAL assay and LPS quantification by ELISA (**K**). All data are plotted as mean ± SEM in male mice, with each symbol on the bar graphs representing a single mouse. *P < 0.05; **P < 0.01; ***P < 0.001; ^#^P < 0.05; ^##^P < 0.01.

To determine how genetic removal of the capacity to generate such HDL would affect liver injury after small bowel resection, we compared liver injury outcomes in *Abca1*^Δ *Vil*^ mice versus control (*Abca1*^fl/fl^) or *Abca1*^Δ *Alb*^ mice 8 weeks after small bowel resection. HDL-C in portal blood was reduced in *Abca1*^Δ *Vil*^ mice (Fig. 5G) that also exhibited higher liver fibrosis and inflammatory parameters (Fig. 5H, I). Transcripts for tumor necrosis factor (*Tnf*), chemokine (C-C motif) ligand 2 (*Ccl2*), and type I collagen (*Col1a1*) were enhanced in livers of *Abca1*^Δ *Vil*^ mice (Fig. 5J), but *Abca1*^Δ *Alb*^ mice did not show altered liver injury compared with controls (Fig. 5H-J). *Abca1*^Δ *Vil*^ mice had elevated LPS activity assessed in the Limulus assay (Fig. 5K), although the absolute amount of LPS in portal plasma was not different from other experimental groups (Fig. 5K). This finding is consistent with the expectations that lower HDL-C in *Abca1*^Δ *Vil*^ mice compared with *Abca1*^fl/fl^ or *Abca1*^Δ *Alb*^ mice would increased LPS activity for a given quantify of LPS, due to reduced HDL-mediated neutralization. Together, these data underscore that intestinal HDL protects against injury in a TLR4-driven model of liver damage.

### Activation of LXR in the intestine increases HDL output and protects against liver injury

Finally, we wondered if we could target selective upregulation of intestinal HDL to promote liver health after bowel resection. The liver X receptors (LXRs) are transcription factors that govern expression of key HDL related genes like *Abca1*. While systemic LXR agonists cause liver steatosis, low-dose LXR agonists such as GW3965, when administered orally, bypasses activation of LXRs in the liver while targeting it in the intestine (*29*), avoiding liver steatosis while reducing atherosclerosis (*30*). After administering GW3965 orally at 1mg/kg, *Abca1* mRNA was elevated ∼4-fold in the ileum of control *Abca1*^fl/fl^ mice, but was abrogated in *Abca1*^Δ *Vil*^ mice; no induction of *Abca1* mRNA was observed in liver in either strain (Fig. 6A). Other LXR downstream genes including *Apoa1* were induced in the ileum, but none in a number of probed LXR-regulated lipogenic genes were altered in the liver in response to oral GW3965 (Fig. S3). Portal venous HDL-C, remaining predominantly in the form of HDL_3_, was enhanced by GW3965 in *Abca1*^fl/fl^ mice but not in *Abca1*^Δ *Vil*^ mice (Fig. 6B). Strikingly, oral GW3965 prominently suppressed appearance of Sirius red^+^ fibrotic area in the liver in response to small bowel resection, while also suppressing infiltration of neutrophil S100A9^+^ cells, plasma AST levels, and induction of inflammatory genes (Fig. 6C-F). The protection of the liver against injury after small bowel resection by oral GW3965 was remarkably abrogated in *Abca1*^Δ *Vil*^ mice (Fig. 6C-F), indicating that the mechanism of action by oral GW3965 depended on raising intestinal HDL.

**Figure 6.**
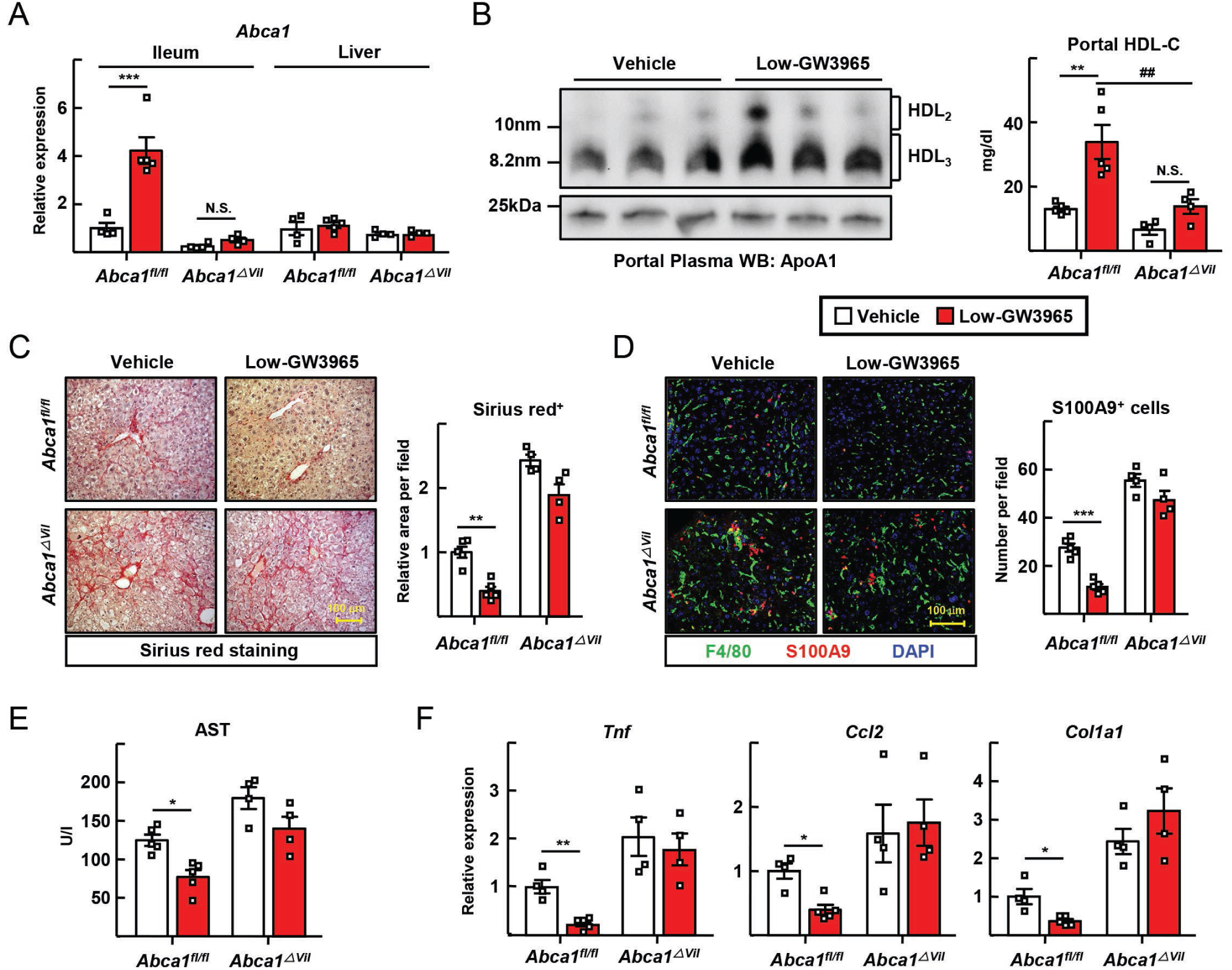
Intestine-specific activation of LXR improves SBR-induced liver injury. Mice received 1 mg/kg/day GW3965 by gavage twice weekly in the last 5 weeks of a 10-week feeding following SBR (Low-GW3965). (**A**) The ileum and liver mRNA transcripts of *Abca1* were analyzed by qRT-PCR. (**B**) Immunoblot for apoA1 run on a nondenaturing gel was conducted on portal vein plasma (left). Portal plasma HDL cholesterol levels (right). (**C**) Sirius red staining of liver sections (left) and relative area per field (right). (**D**) F4/80^+^ macrophages and S100A9^+^ neutrophils were visualized by immunofluorescence in liver sections (left) and cell numbers per field counted (right). (**E**) Plasma AST levels. (**F**) Hepatic mRNA transcripts. All data are plotted as mean ± SEM in male mice, with each symbol on the bar graphs representing a single mouse. *P < 0.05; **P < 0.01; ***P < 0.001; ^##^P < 0.01.

## Discussion

Intestinal villi are highly organized structures that permit intestinal epithelial cells to deposit cargo, especially absorbed nutrients, basolaterally so that it can be readily taken into the bloodstream and then to the portal vein, or to lymphatics if molecules are too big for entry into fenestrated blood vessels (*16, 31*). Intestinal epithelial cells also produce HDL particles (*11*), but neither the fate nor the function of intestinal HDL has been clear. We show that the intestine produces the smaller form of HDL, called HDL_3_, enriched in anti-inflammatory proteins like PON1. This HDL does not get routed to lymphatics, like it is in other tissues (*16*), but is shuttled to the portal vein to directly access liver. Functionally, this HDL binds and neutralizes LPS, much more so than HDL_2_. Thus, both the trafficking fate and the function of intestinal HDL are coordinated to specifically serve a role in host immune protection by the downmodulation of liver injury through, at least in part, the neutralization of LPS and reduction of TLR4 signalling in the liver. The intestinal epithelial location for HDL production can support local, immediate capture of LPS before it gains access to and promotes inflammation in underlying host tissue, in the portal vein itself, and ultimately the liver. This mechanism is likely most important when the intestine has been subjected to a homeostasis-altering insult like the dramatic surgical resection and adaptation we study here. From this perspective, it is also fitting that the highest expression of apoA1 and ABCA1, critical for HDL synthesis, coincides with the part of the small intestine, the ileum, with the richest microbiome and consequently the highest accumulation of LPS. Furthermore, this model provides a framework to consider how loss in the expression of HDL components like apoA1 in Crohn’s disease of the ileum (*32*), a major form of inflammatory bowel disease, might enhance inflammatory injury.

Our study brings forward some as yet unanswered questions in basic science that will be important to pursue in future studies. One of these questions is how LPS and LBP interact with HDL-C and why HDL_3_ is more likely than HDL_2_ to bind to LPS and neutralize it. Related is the question of whether the ability to neutralize LPS is linked to other anti-inflammatory properties of the HDL particle or involves proteins on HDL_3_ like PON-1. In other clinical settings, especially cardiovascular disease and sepsis, HDL_3_ levels, not HDL_2_, correlate with better health outcomes (*33, 34*).

Another mystery is why the vast majority of portal venous HDL-C derives from the intestine, a finding that first became apparent when we found little HDL-C in the portal vein of mice selectively lacking intestine-derived HDL and was further supported a series of experiments in which we followed a photoconvertible version of HDL that we earlier engineered for fate mapping of HDL trafficking. A priori one might have thought that even if intestine-derived HDL had a portal venous fate rather than a lymphatic one, intestinal HDL would mix with liver-derived HDL arriving from the systemic blood supply to the intestine, such that the portal venous blood HDL would arise from both portal and systemic sources. However, we find that, while systemic HDL clearly arrives to the gut or mesentery, because it efficiently enters the intestine-draining mesenteric lymphatics after originating from a distal systemic site like the skin, it poorly enters the portal vein. These data suggest existence of unknown trafficking steps, possibly wherein HDL-C from the systemic circulation becomes accessible to the interstitium of the mesentery, muscularis, submucosa, or lamina propria before being picked up by the lymphatics. This hypothetical accessibility of HDL to the interstitium is not with possible parallels, as it has been shown that at least some fluorescent tracers readily leak out from incoming systemic blood into the lamina propria (*19*). HDL_2_ may be too large to enter the fenestrated blood vessels that drain to the portal blood, such that only intestine-derived HDL_3_ gains efficient access. Consistent with this possibility, mesenteric lymph is relatively deficient in the smaller HDL_3_ particles while relatively enriched in HDL_2_ (*35*).

These studies have strong translational potential. In human subjects, as in mice, the portal blood was enriched in HDL_3_ and potently suppressed Kupffer cell activation in the presence of LPS. We also utilized a mouse model of liver injury, that of small bowel resection, that models a surgical intervention applied in humans that leads to liver failure in many patients. From a therapeutic perspective, oral delivery of LXR agonists proved effective in protecting the liver by upregulating HDL within the intestine. LXR agonists have failed to find utility in the clinic to date, but orally restricted LXR agonists, especially those like GW3965 designed to remain with the intestine itself (*36*) remain promising (*37*). Our findings highlight an entirely new application, treatment of short bowel syndrome, for their potential clinical use. Furthermore, HDL-mediated protection from liver injury in this model is likely applicable to other causes of liver injury linked to changes in the microbiome or intestinal permeability. If suitable LXR agonists cannot be developed for application in humans, other approaches to elevate intestinal HDL should be explored, as doing so seems likely to improve health in the gut-portal-liver axis.

## Acknowledgements

We are grateful to Mary Sorci-Thomas (Medical College of Wisconsin) for critical reading of the manuscript and to Mary Wohltmann for expert mouse colony maintenance. This work was supported primarily by NIH R01 DK119147 to GJR and BWW, with additional support from DP1DK1109668 and AI0499653 to GJR, with the latter including a Primary Caregiver Supplement to support career development of EJO. EJO also received support from T32 DK077653. LH is supported by an American Heart Association Career Develop ment Award (AHA: 18CDA34110273). RSC is supported by the Lawrence C. Pakula, MD IBD Research Fellowship. Histology was carried out using core facilities supported by the Digestive Diseases Research Core Center of Washington University, P30 DK052574. Electron microscopy was performed at the Washington University Center for Cellular Imaging (WUCCI), which is funded in part by the Children’s Discovery Institute of Washington University and St. Louis Children’s Hospital (CDI-CORE-2015-505 and CDI-CORE-2019-813) and the Foundation for Barnes-Jewish Hospital (3770).

## Author Contributions

Y.H.H, E.J.O, G.J.R designed experiments; Y.H.H, E.J.O, L.H.H, R.S.C conducted experiments and developed techniques; Y.H.H analyzed data; Y.H.H, G.J.R wrote the manuscript; L.H.H, R.S.C, B.W.W edited the manuscript; G.J.R, B.W.W obtained funding and compliance approvals.

## Supplementary Materials

### Materials and Methods

#### Mouse colony and surgical or treatments

C57BL/6 WT, TLR4^−/−^ (B6.B10ScN-Tlr4^lps-del^/JthJ), *Villin*-Cre (JAX #002216), *Albumin*-Cre (B6.FVB(129)-Tg(Alb1-cre)1Dlr/J), *Abca1*^*fl/fl*^ (B6.129S6-Abca1^tm1Jp^/J) mice (7–10 weeks of age) were purchased from the Jackson Laboratories and housed in a specific-pathogen-free room at 22– 24°C and 50–60% humidity with a 12 h light/dark cycle. were purchased from Jackson Laboratories. We previously generated and described PGA^KI/+^ mice. All experiments were performed in a blinded and randomized fashion. Mice were housed on a 12-hour light-dark cycle in a temperature controlled, specific pathogen-free unit with food and water provided ad libitum. The study was approved by the Washington University Animal Studies Committee (Protocols 20170154, 20170252 and 20-0032) in accordance with the National Institutes of Health laboratory animal care and use guidelines.

For small bowel resection experiments, mice underwent a 50% proximal (jejunal) bowel resection, 75% proximal bowel resection, or sham control operation (bowel transection with reanastomosis alone), as previously described (*2*). In brief, through a midline laparotomy, the small bowel was exteriorized and transected 1 to 2 cm distal from the ligament of Treitz and approximately 12 cm (for 50% resection) or 6 cm (for 75% resection) proximal to the ileocecal junction. For sham operations, a transection 12 cm proximal to the ileocecal junction with immediate re-anastomosis were performed. For distal 50% SBR, the ileum (last 12 cm of small bowel) was removed with an anastamosis of the jejunum to a small cuff of small bowel on the cecum. All anastomoses were hand-sewn end-to-end with interrupted 9-0 nylon sutures. Post-operative care included housing in an incubator for temperature stability and 24 h fasting before starting a liquid diet (PMI Micro-Stabilized Rodent Liquid Diet LD 101; TestDiet), which they were maintained on for 8-12 weeks until euthanasia.

For bone marrow transplants, WT recipient mice at 8 weeks of age received whole-body irradiation at a dose of 11 Gy, and then were intravenously injected with 5 ⨯ 10^6^ bone marrow cells from wild type or TLR4^−/−^ donor mice. After 4 weeks, short bowel resectionswere conducted.

For LXR agonist experiments, GW3965 (Sigma-Aldrich, #G6295) was suspended in 0.5% carboxymethyl cellulose and was orally administered twice weekly at 1 mg/kg/day for the last 5 weeks in the 10-week period following intestinal resection. The different experimental groups of mice maintained a similar body weight during liquid diet feeding and/or drug treatment.

#### Immunostaining and confocal microscopy

Left lobes of liver tissues and small intestines were excised and fixed in 4% paraformaldehyde (Santa Cruz Biotechnology) overnight at 4° C. 10-μm paraffin-embedded sections were treated for antigen retrieval with Diva Decloaker (Biocare Medical; #DV2004) in boiling, pressurized chamber for 15 min. Sections were blocked with 5% donkey serum, 1% bovine serum albumin (BSA) (Sigma-Aldrich) and 0.03% Triton-X (Plusone, #17-1315-01) in PBS for 1 hr, then incubated in with rat anti-F4/80 (Abcam, #ab6640), goat anti-S100A9 (R&D Systems, #AF2065), rabbit anti-von Willebrand Factor (DAKO, #a0082), or rat anti-PV1 (BD pharmingen, #550563) at 4°C overnight. Primary antibodies were detected using Cy3- or Cy5-conjugated secondary antibodies (Jackson ImmunoResearch). The stained sections were imaged using an SP8 confocal microscope (Leica) and images were processed with Imaris software (Bitplane). Ten fields were quantified and averaged for each sample, with cell counts per image acquired at 200X magnification using Image J software (NIH). All slides were analyzed by in blinded and randomized fashion.

#### Quantitative real-time polymerase chain reaction (qRT-PCR)

Total RNA from tissues or cells was isolated by using RNeasy Mini or Micro kits according to the manufacturer’s protocol (Qiagen). cDNA was synthesized using high-capacity cDNA reverse transcription kit (Applied Biosystems, #4368814). qRT-PCR experiments were performed using ABI StepOnePlus Real-Time PCR machine with specific primers (Applied Biosystems). Primer sequences are available upon request. The relative transcriptional expression of target genes was evaluated by the equation 2^−ΔCt^ (ΔCt = Ct of target gene minus Ct of 18S rRNA). Relative transcriptions in the mRNA level of each genes were calculated with a the mean of the control group set as 1.

#### Intestinal permeability assay

After fasting 4 h, 200 mg/kg of 4 kDa FITC-dextran (Sigma-Aldrich) was administered by gavage. 2h later, blood for the preparation of plasma was collected from the tail vein, and the fluorescence intensity of the samples and standards were read at excitation 485 nm / emission 525 nm using the Cytation 5 Cell Imaging Multi-Mode Reader (BioTek).

#### Photoactivation of PGA1 ^KI/+^ mice

To track the trafficking fate of photoconvertible HDL from skin, 8-10 wk old PGA1^KI/+^ mice were anesthetized and a region of shaved skin was photoconverted using a SOKY, Violet 405nm 500 mW (FDA), PL-405-500B laser, as described previously (*1*). For photoactivation of at the lumen of the small intestine of anesthetized mice, we stretched the mesentery and intestine over the solid surface of a petri dish, located the region of interest and surgically clipped the bowel just enough so that we could thread into the lumen a fibro-optic endoscopic laser (Laserland, Violet 405nm 100 mW) to photoactivate enterocytes. 3 areas were activated for one data point, with in each case, the laser being held on 10 seconds, with 20 seconds off, cycling for 1 min 10 seconds to achieve 3 exposures per location. For photoactivation of the exterior of the small intestine, the Laserland, Violet 405nm 100mW 5V laser was used to activate area of 14.668 mm^2^ of gut with a similar on/off cycle as for the intestinal lumen.

#### Collection of blood and lymph

The portal blood was collected with a 33-gauge needle, 50 µl per mouse; systemic blood was collected from the inferior vena cava using a 26-gauge needle in EDTA-containing tubes, or mesenteric lymph fluid was collected, and fluorescence intensity of plasma or lymph fluid was measured using the Cytation 5 Cell Imaging Multi-Mode Reader (BioTek). Mesenteric lymphatic cannulations were accomplished under general anesthesia using an operating microscope. A midline laparotomy with an extension to a left subcostal incision was made and the intestine was mobilized to expose the mesenteric lymphatic duct proximal to the cisterna chyli. A small incision was made on the mesenteric lymphatic duct and gently cannulated using polyethylene tubing (ID .28mm OD .61mm, Intramedic, Sparks, MD). At the completion of the collection, the mouse was euthanized.

#### Preparation of HDL fractions

Human and mouse plasma collected from the portal vein or peripheral vein (inferior vena cava for mouse, antecubital vein for human) were collected and ultracentrifugated for overnight at 45,000 rpm, 4°C with PBS buffer containing EDTA, sodium chloride, and potassium phosphate dibasic. Chylomicron and VLDL fractions floating first to the top were carefully removed. Then the density of the solution was adjusted to 1.063 g/ml using KBr (Sigma-Aldrich; #221864) buffer and subjected another overnight centrifugation using the same parameters as above. The LDL fraction now emerging as the top layer was next removed and, finally, the HDL fraction was collected after another overnight centrifugation at the same speed in 1.21 g/ml KBr buffer. Isolated HDL fractions were dialyzed using Slide-A-Lyzer Dialysis cassette kit (Thermo Fisher) with PBS solutions containing sodium chloride, Tris and EDTA for 4 hr at 4°C to remove KBr. Human HDL_2_ (1.063 – 1.124 g/ml) and HDL_3_ (1.12 – 1.21 g/ml) fractions were obtained from GenWay Biotech.

#### FPLC and measurement of HDL cholesterol

Fifty microliters of blood was collected in Eppendorf tubes containing 10 μl 0.5 mM EDTA and then centrifuged for 6 min at 6,000 rpm. For cholesterol distribution of total lipoproteins, plasma was prepared and 100-200 μl of the plasma was flowed over a Superose 6 10/300GL gel filtration column (GE Healthcare) to separate the different classes of lipoproteins. Cholesterol in each fraction was measured by an enzymatic assay kit (Wako Diagnostics Cholesterol E; #439-17501). HDL cholesterol assay kit (Cell Biolabs; #STA-394) was used to measure HDL-specific cholesterol levels (HDL-C).

#### Immunoblots

Western blotting was performed using rabbit anti-mouse apoA1 (Meridian Life Sciences), rabbit anti-human apoA1 (Millipore, #MAB011), rabbit anti-human ABCA1 (Novus Biologicals, #NB400-105), mouse anti-PON1 (Abcam, #ab24261), rabbit anti-ApoB (Proteintech, #20578-1), or rabbit anti-albumin (Proteintech, #16475-1) antibodies, as described previously (*1*). The HDL fractions were loaded to achieve the same protein concentration per lane, and plasma loaded without dilution. For native gels, the samples were diluted in 2X native sample buffer (Bio-rad) and run on 4-20% Mini-PROTEAN Tris-glycine gels (Bio-rad) with Tris-glycine running buffer. For denaturing gels, the samples were diluted in 2X Tris-Glycine-SDS sample buffer (EZ Bioresearch) and heated at 95°C for 10 min. The samples were loaded onto 4-20% Mini-PROTEAN gels and run with Tris-glycine-SDS running buffer. The separated proteins were transferred to 0.45 μm PVDF membrane (Milipore,#IPVH00010) with Tris-glycine transfer buffer for 2 hrs at 20 V. Membranes were blocked with 5% nonfat dry skim milk (Bio-rad, #170-6404) for 1 hr and primary antibodies were incubated overnight at 4°C. After incubation with HRP-conjugated secondary antibodies, signal detection was done using Clarity Western ECL solution (Bio-rad).

#### Electron Microscopy of HDL particles

The isolated HDL fractions were diluted to 15 μg/ml total protein concentration and negatively stained with 1% uranium acetate. The samples were deposited on carbon-coated 200 mesh copper grids (Electron Microscopy sciences). Images were acquired with a TEM (JEOL JEM-1400^Plus^) at 120KeV and 80,000x or 150,000x magnifications. The diameter of HDL particles was measured using Image J software.

#### Isolation or culture of liver immune cells and macrophages

For quantification of neutrophils, monocyte-derived macrophages and Kupffer cells in livers of mice subjected to short bowel resection or sham surgery, livers were collected and homogenized in Hank’s buffered saline containing 1.49 mg/ml collagnase type IV (Sigma-aldrich; #C5139) and dissociated using the gentleMACS Octo Dissociator (Miltenyl Biotec). After a low speed centriguation, the supernatant containing nonparenchymal cells was separated using 33% Percoll (GE Healthcare).

For Kupffer cell isolation and culture, the livers of 7–10-week-old male C57BL/6 mice were perfused via the inferior vena cava with collagenase type IV solution as described previously (*3*).Cell suspensions were centrifuged in 50%/25% Percoll (GE Healthcare). The layer containing liver macrophages was plated in RPMI-1640 (Hyclone) containing 10% fetal bovine serum (FBS). After 2 h culture to allow for cell attachment, the cell medium was changed to “vehicle” medium, which was serum-free RPMI-1640 containing 1 μg/ml recombinant LBP (R&D systems; #6635-LP) for 3 h of culture. HDL preparations as described were added, or not, to these cultures with LPS (Sigma-Aldrich; #L2630) presence.

#### Flow cytometry

Isolated liver immune cells and cultured liver macrophages were collected and counted in an automated cell counters (Cellometer Auto X4, Nexelcom Bioscience) after staining for acridine orange (Sigma-aldrich). FACS antibodies including BUV396-anti-CD45 (BD bioscience; # 563791), FITC-anti-Ly6G (Biolegend; #127605), APC/Cy7-anti-F4/80 (Biolegend; #123117), PerCP/Cy5.5-anti-Ly6C (Biolegend; #128011), PE/Cy7-anti-CD31 (Biolegend; #102417), Alexa488-anti-iNOS (Thermo Fisher; #53-5920-82), APC-anti-CD11b (Thermo Fisher; 17-0112-82), PE-anti-Tim4 (Thermo Fisher; 12-5866-82), or goat anti-Clec4f (R&D systems; #AF2784) were incubated with FACS buffer (5% fetal bovine serum, 5 uM EDTA, and sodium azide in PBS) on ice for 30 min. For intracellular staining, we used IC fixation and permeabilization buffers as per manufacturer’s guidance (Thermo ebiosciences). After washing and resuspension, cells were analyzed on a BD FACS Symphony machine and analyzed by FlowJo software (BD biosciences).

#### ELISA

ELISA kits were used according to manufacturer’s protocol including AST Activity Assay Kit (Sigma-Aldrich; MAK055). TNFα (Siglma-Aldrich; RAB0477), or CCL2 ELISA (Sigma-Aldrich; #RAB0055). Limulus amebocyte lysate (LAL) endotoxin activity was measured using the Pierce LAL Chromogenic Endotoxin Quantitation Kit (Thermo Fisher; #88282). LPS quantification by ELISA used LPS ELISA kit from MyBiosource (MBS700021). For sandwich ELISA to analyze LPS-HDL binding, we used high-binding clear polystyrenes microplates (R&D systems, DY990), and purified HDL was immobilized for 2 h on these plates at 37°C at 10 μg/ml. After washing, plates were blocked with 1% BSA for 1 h. Then Biotin-LPS (InvivoGen; #tlrl-lpsbiot) was pre-incubated with or without 1 μg/ml recombinant LBP (R&D Systems) for 1 h and incubated in plates for 30 min. Streptavidin peroxidase (R&D Systems) was added, followed by diaminobenzidine substrate (Abcam), for colorimetric reactions. All colorimetric and fluorometric absorbances were detected using the Cytation 5 Cell Imaging Multi-Mode Reader (BioTek).

#### Human Studies

Human portal and peripheral systemic blood was acquired from adult patients undergoing open surgical procedures where the surgical team deemed that the portable vein was safely accessible. Blood were collected in EDTA-containing tubes, spun down for immediate analysis of plasma for HDL. Patients, 3 males and 3 females, ranged in age from 54 to 80; 4 were receiving Whipple pancreatic surgery, one gastric bypass, and the other orthotopic liver transplant. All human studies were approved by the Human Research Protection Office at Washington University, IRB protocol # 2019101009 (Randolph, G. J., PI).

#### Statistics

All values are expressed as the mean ± SEM. Statistical differences were determined by the unpaired two-tailed Student’s t test for simple comparisons or one-way ANOVA with Tukey’s post hoc test for multiple comparisons for 3 or more groups. *P* < 0.05 was determined as significantly different. Statistical differences were analyzed by using GraphPad Prism Software Version 8.0 (GraphPad Software).

## Supplementary Figures and Legends

**Figure S1.**
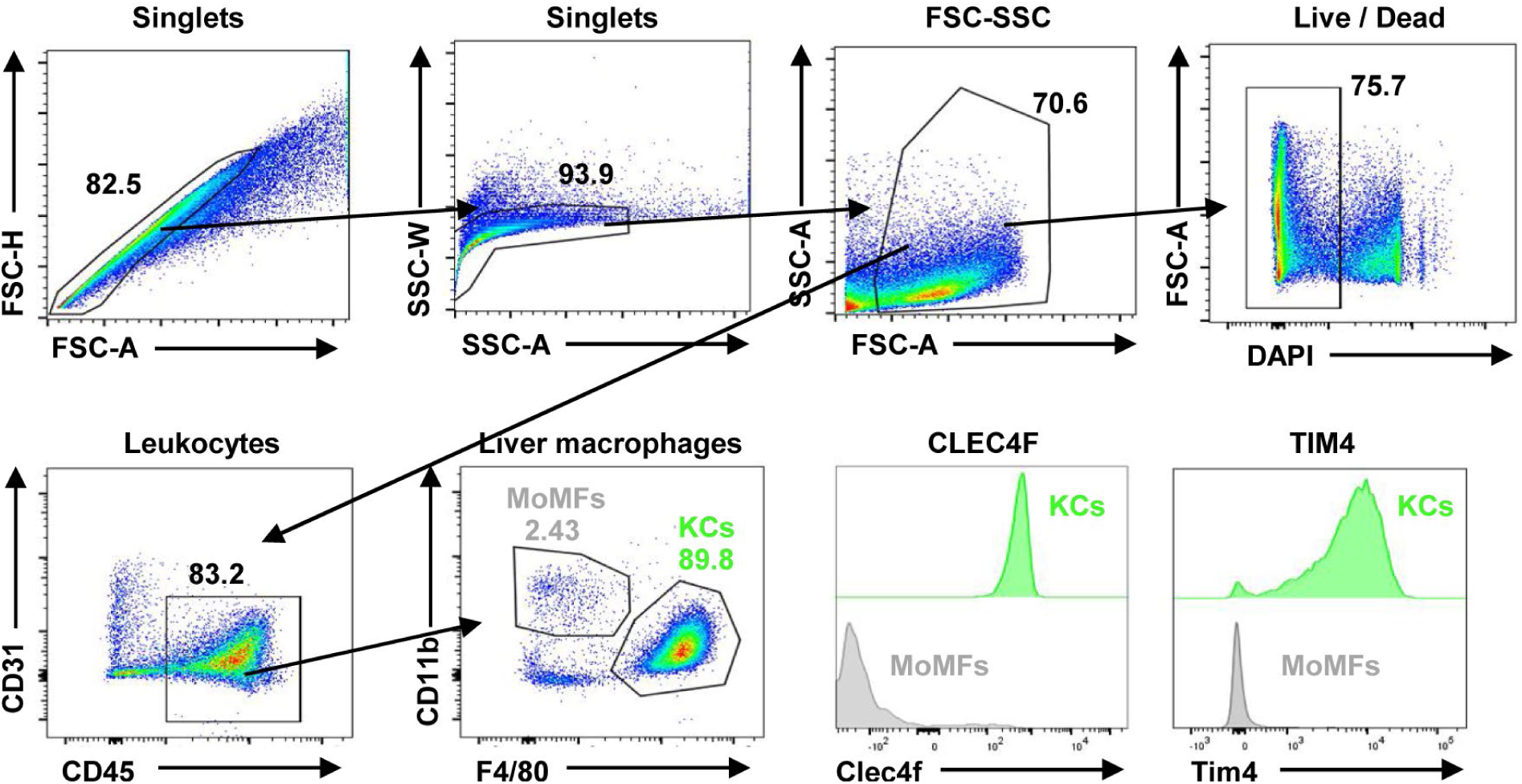
Flow cytometric analysis and gating strategy of liver macrophage Kupffer cells. Gating strategy and representative flow dots of liver macrophages, Kupffer cells (KCs), and monocyte-derived macrophages (MoMFs). Residential Kupffer cells were over approximately 80% purity among liver leukocytes after isolations.

**Figure S2.**
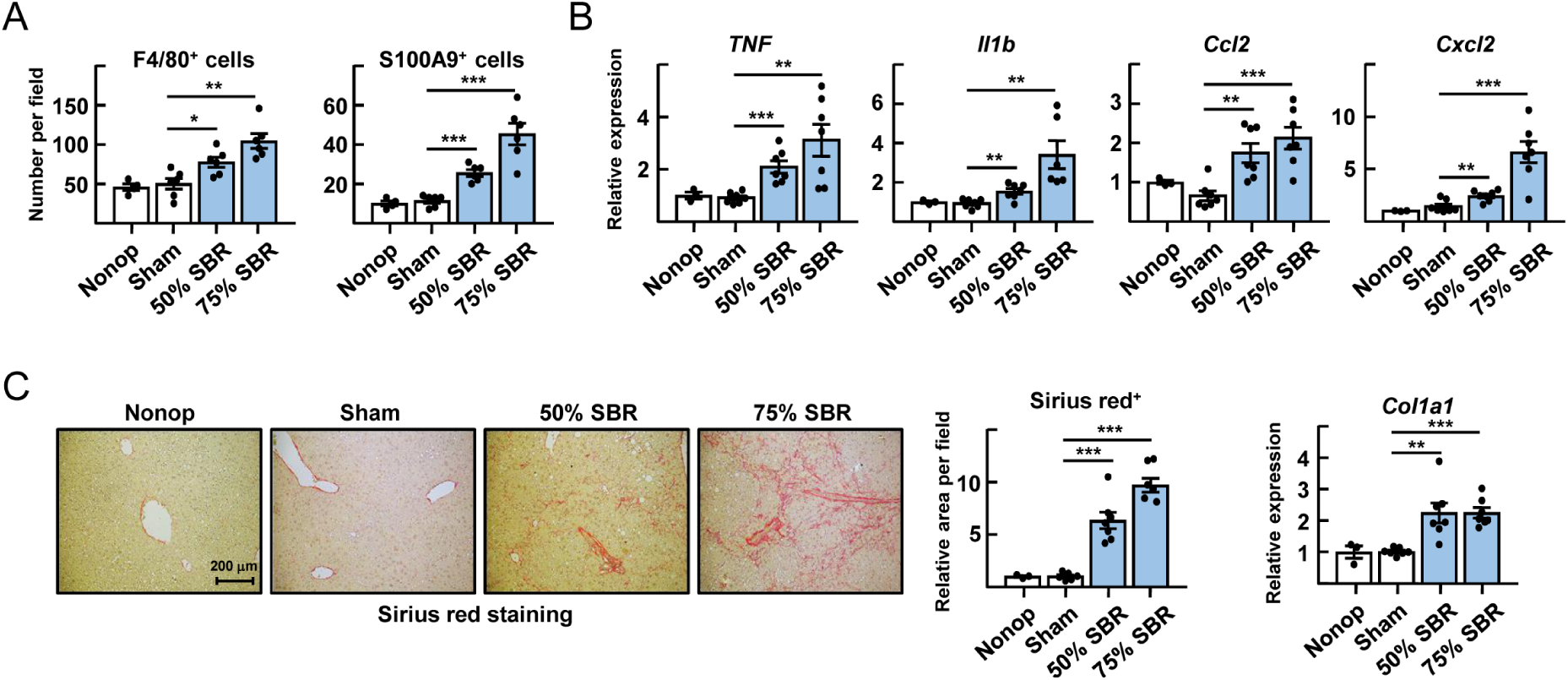
Short Bowel resection induces liver inflammation and fibrosis. Small bowel resection operations were carried out on WT mice. Non-operated (Nonop) (n=4), sham (n=8), 50% SBR (n=8), and 75% SBR (n=8) mice were euthanized 12 weeks later. (**A**) The cell numbers of F4/80^+^ macrophages and S100A9^+^ neutrophils per X200 fields were counted. (**B**) The hepatic mRNA levels of inflammatory genes were analyzed by qRT-PCR. (**C**) The representative images of sirius red staining of liver sections (left). The hepatic mRNA levels of *Col1a1* were analyzed by qRT-PCR (right). All data are represent as mean ± SEM. *P < 0.05; **P < 0.01; ***P < 0.001.

**Figure S3.**
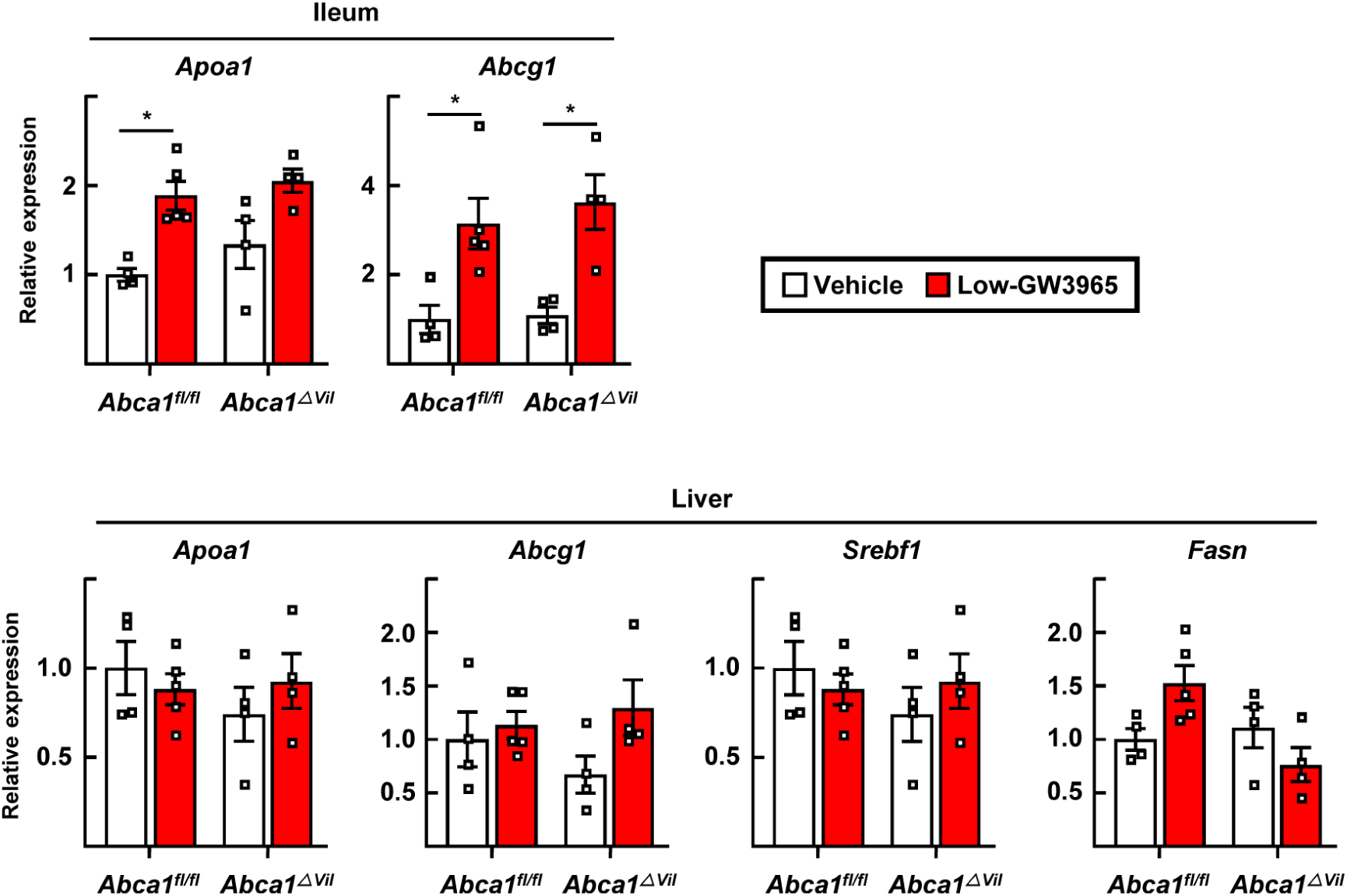
Oral administrations of low doses GW3965 only influences intestines, not livers. Mice received 1 mg/kg/day GW3965 by gavage twice weekly in the last 5 weeks of a 10-week feeding following SBR (Low-GW3965). The ileum and liver mRNA transcripts of *lxr* downstreme genes were analyzed by qRT-PCR. All data are represent as mean ± SEM. *P < 0.05; **P < 0.01; ***P < 0.001.

